# POU2AF2/C11orf53 functions as a co-activator of POU2F3 by maintaining chromatin accessibility and enhancer activity

**DOI:** 10.1101/2022.03.17.484753

**Authors:** Aileen Patricia Szczepanski, Natsumi Tsuboyama, Zibo Zhao, Lu Wang

## Abstract

Small cell lung cancer (SCLC), accounting for around 13% of all lung cancers, often results in rapid tumor growth, early metastasis, and acquired therapeutic resistance. The POU class 2 homeobox 3 (POU2F3) is a master regulator of tuft cell identity and defines a subtype of SCLC tumors (SCLC-P subtype) that lack the expression of neuroendocrine (NE) markers. Here, we have identified a previously uncharacterized protein, C11orf53, which is co-expressed with POU2F3 in both SCLC cell lines and patient samples, defining the same SCLC-P subtype. C11orf53 is an essential gene for the cell survivability of SCLC-P subtype, based on genome-wide CRISPR screening and genetic depletion of C11orf53 or POU2F3, inducing apoptosis in different SCLC-P subtype cell lines. Furthermore, our biochemical and genome-wide studies have demonstrated that C11orf53 is a novel chromatin-bound protein that occupies broad active enhancer domains and regulates gene expression at active enhancers. Mechanistically, C11orf53 directly interacts with POU2F3 at the POU domain, and is recruited to chromatin by POU2F3. Consequently, depletion of *C11orf53* dramatically reduced enhancer H3K27ac levels and chromatin accessibility, resulting in a reduction of the expression of POU2F3-dependent tuft cell markers. Based on the expression pattern and molecular function of C11orf53, we have renamed it as “POU Class 2 Homeobox Associating Factor 2” (POU2AF2). In summary, our study has identified a new co-activator of POU2 family transcription factors, and sheds light on the therapeutic potential of targeting POU2AF2/POU2F3 heterodimer in human SCLC.

## Introduction

Lung cancer is one of the leading causes of cancer death world-wide^1^. Specifically, small cell lung carcinoma (SCLC) is a highly aggressive and lethal form of malignancy that develops within tissues of the lungs and is typically diagnosed at later stages when it has already metastasized^2-4^. Although SCLC consists of approximately 13% of all lung cancer cases, it has one of the lowest 5-year survival rates and a very poor prognosis^5^.

Recent studies have made advancements in the classification of SCLC defined by relative expression of four major molecular subtypes of key lineage-specific transcription and co-transcription regulators: achaete-scute homolog 1 (ASCL1; SCLC-A), neurogenic differentiation factor 1 (NEUROD1; SCLC-N), yes-associated protein 1 (YAP1; SCLC-Y), and POU class 2 homeobox 3 (POU2F3; SCLC-P)^6^. Analysis of the morphological features have further indicated that SCLC can be classified into neuroendocrine (NE)-high (SCLC-A and -N) or -low (SCLC-Y and SCLC-P) subtypes based on the expression pattern of different NE markers and the diversity in genetic alterations, growth properties, and immune infiltration^7-10^.

Previously, we have identified an essential epigenetic co-regulator and biomarker, additional sex combs-like protein 3 (ASXL3), which is associated with SCLC-A molecular subtype. ASXL3 functions as a scaffold protein that links histone H2A deubiquitinase BAP1 and the Bromodomain-containing protein BRD4 at active enhancer and drives lineage-specific transcriptional programming^11^. Therefore, our study suggested it is necessary to globally identify and study these subtype-specific essential factors to gain a more in-depth understanding of underlying mechanisms regulating SCLC tumorigenesis which may potentially assist in treatment decisions for SCLC and create a much-needed individualized strategic approach.

For instance, the SCLC-P subtype, which is defined by the expression of a master transcription factor, POU2F3, is known to be a variant form of SCLC that lacks NE features^12^. However, identifying other contributing factors in the progression of SCLC-P, specifically a co-factor of POU2F3 function, has yet to be determined. By our current dependency data analysis, and subsequent biochemical and genetic studies, we have identified a previously uncharacterized gene, *C11orf53*, as a co-dependent gene of POU2F3. In these studies, we have comprehensively uncovered an essential role of C11orf53 as a co-activator of POU2F3 in regulatory gene expression to establish the cell identity of SCLC-P subtype.

## Results

### Landscape of SCLC subtype-specific dependency

Global gene expression profiling has been widely used to define molecular subtypes in human cancers^13,14^, including lung cancer^6^. However, not all highly expressed genes are essential for tumor cell growth or cell viability. Therefore, to identify the functional subtype-specific dependent factor within the four SCLC molecular subtypes, we used CCLE SCLC cell lines^15^ encompassing all four subtypes^6^ defined previously and explored the genes that are most selectively essential to each individual subtype compared to the other classifications. Following the criteria for target gene effect score (≤ −0.5; median difference < 0.2 versus median value of the average of all other groups), we identified 48 genes for SCLC-A, 88 genes for SCLC-N, 66 genes for SCLC-P, and 177 genes for SCLC-Y subtypes which are defined as being selectively essential in each SCLC subtype (Fig. 1A). The expression levels of these genes were compared with no global up- or down-regulation in each subgroup (Fig. S1A). The essential genes in each subtype are enriched in distinct pathways (Fig. 1B), suggesting that each subtype may depend on a unique set of signaling pathways for cell survivability and proliferation. Next, we highlighted the top genes essential or redundant in each subtype (Fig. 1C), revealing the non-overlapping essentiality profiles in these subtypes. The gene effect scores of the four markers–*ASCL1, NEUROD1, POU2F3*, and *YAP1*–showed the selective dependency (Fig. 1D), which is consistent with previous studies for the function of each gene.

**Figure 1.**
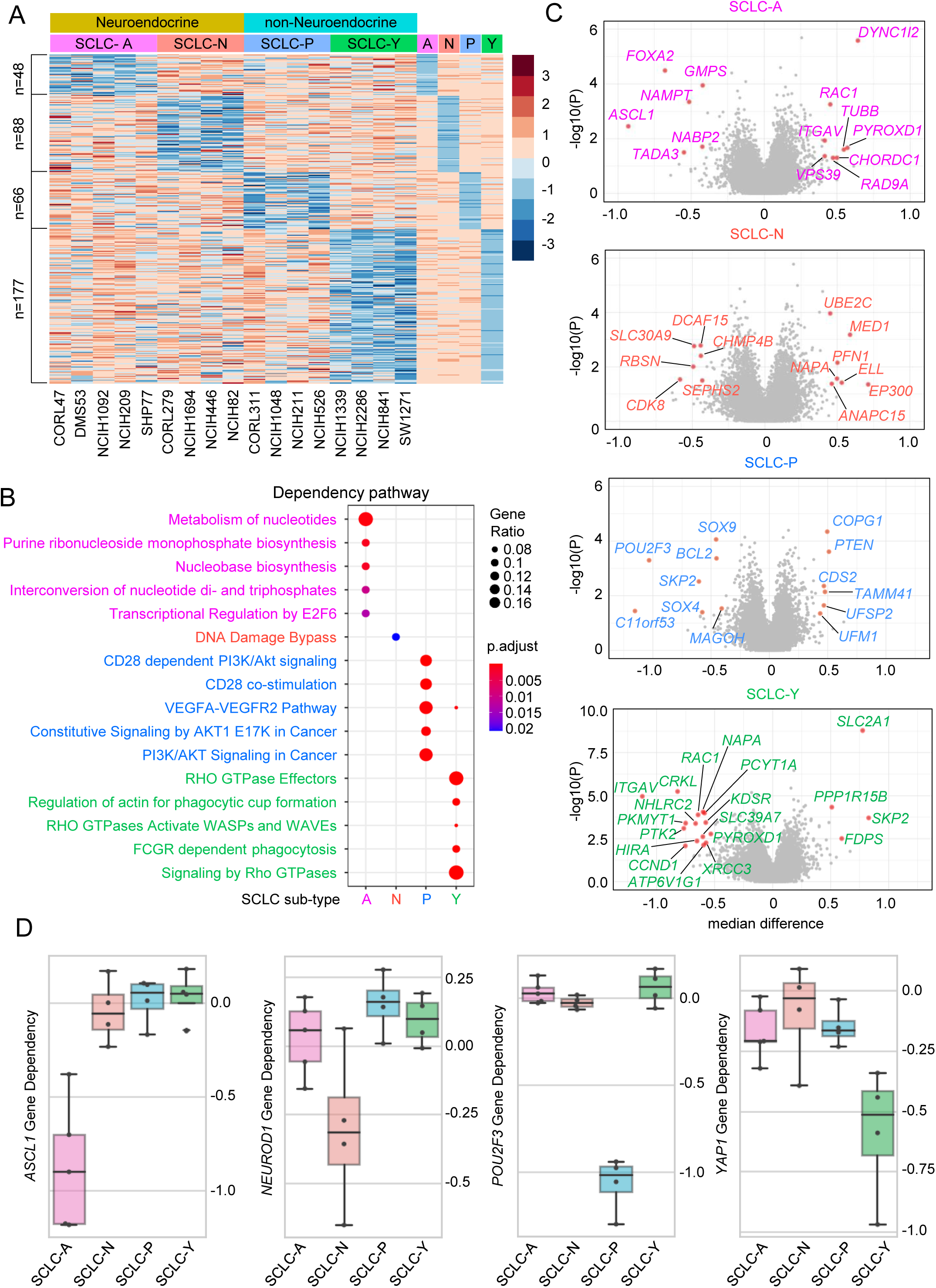
Landscape of SCLC subtype-specific dependency. A) Gene effect scores of all genes were retrieved from DepMap Public 21Q3 datasets for the following 17 SCLC cell lines: A-subtype, CORL47, DMS53, NCI-H1092, NCI-H209, SHP77; N-subtype, CORL279, NCI-H1694, NCI-H446, NCI-H82; P-subtype, CORL311, NCI-H1048, NCI-H211, NCI-H526; Y-subtype, NCI-H1339, NCI-H2286, NCI-H841, SW1271. A- and N-subtypes were classified as neuroendocrine (NE) and P- and Y-subtypes were classified as non-NE groups. For each subtype, the median gene effect scores of each subtype were calculated and the Z-score heatmap showed genes selectively essential in each group with the criteria that target gene effect score is <= −0.5 and the median difference is 0.2 less than the median value of the average of all other groups. B) Metascape pathway analysis of the essential genes in each subtype as identified above. C) Volcano plots of the gene essentiality in the four subtypes. X-axis is the median difference between the indicated group and others, and y-axis is negative log10 p-value calculated with T-test for the means of two independent samples of scores. Highlighted genes have p < 0.05 and median difference > 0.4 for SCLC-A, SCLC-N, and SCLC-P; p < 0.01, median difference > 0.5 for SCLC-Y subtypes. D) Box plots showed the ASCL1, NEUROD1, POU2F3 and YAP1 gene dependency in each subtype.

### Identification of C11orf53 as a new marker for SCLC-P subtype

Previous studies have identified that the transcription factor POU2F3 is a master regulator of a tuft cell-like variant of small cell lung cancer^12^. Therefore, POU2F3 has been widely used to define the SCLC-P subtype. Interestingly, based on our dependency data analysis in Figure 1A, we noted that the gene effect scores related to a gene called chromosome 11 open reading frame 53 (*C11orf53*) (Fig. S2A) is selectively lower in rank (1/66) versus POU2F3 (2/66) in SCLC-P (Fig. 2A). *C11orf53* is an uncharacterized gene that is an evolutionarily conserved sequence from zebrafish to human (Fig. S2B, C) and is expressed at high levels in all four SCLC-P cell lines examined (Fig. S2D).

**Figure 2.**
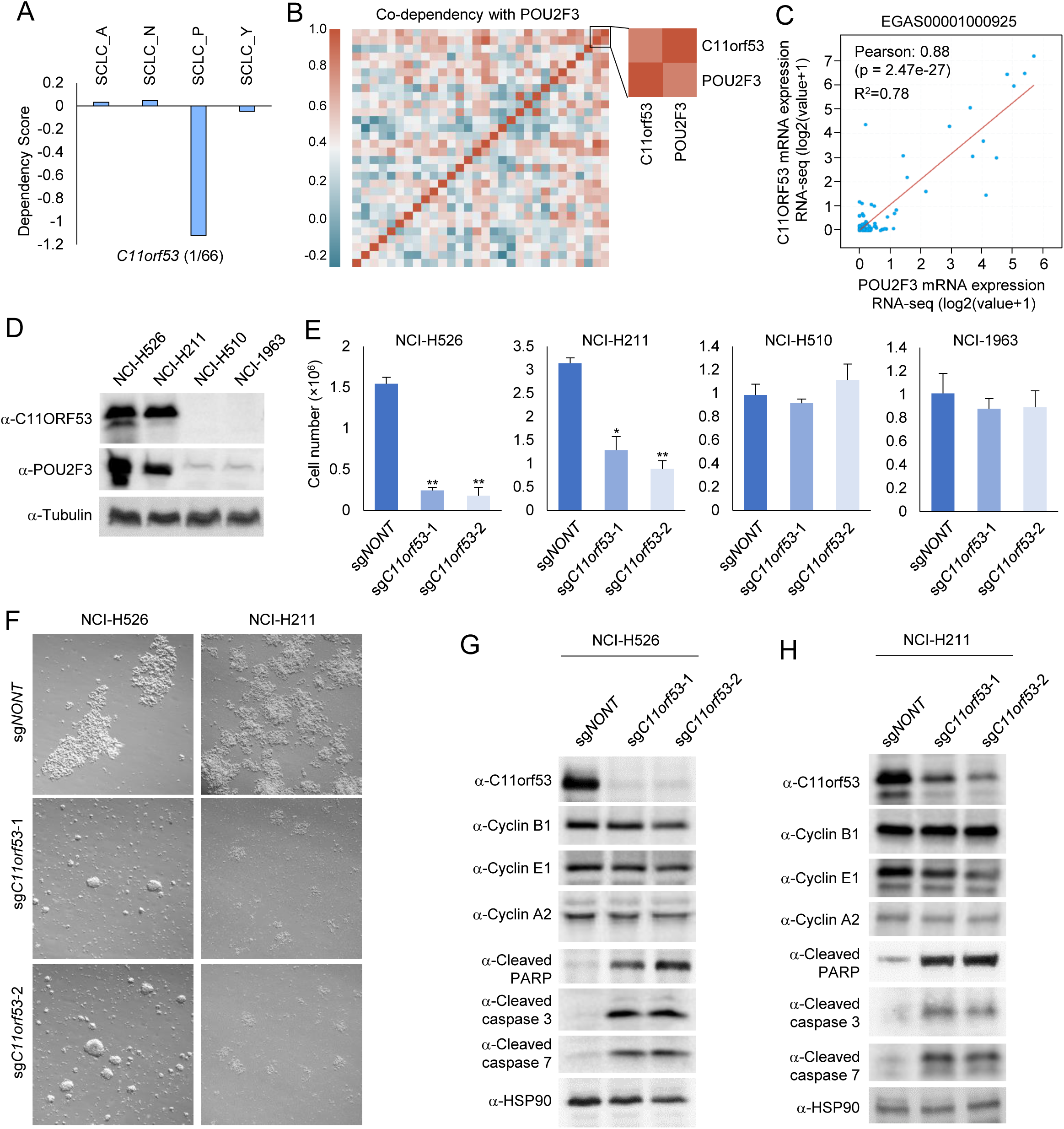
Identification of C11orf53 as a new marker for SCLC-P subtype. A) Box plots showed the C11orf53 gene dependency in each subtype. B) Co-dependency matrix showing the pearson correlation coefficient values of the top 30 genes selectively essential in SCLC-P subtype. C) The scatter plot shows the correlation between POU2F3 and C11orf53 mRNA levels in 110 patient samples (EGAS00001000925). D) The protein levels of C11orf53 was determined by western blot in four different SCLC cell lines, n=2. E) Four different SCLC cell line NCI-H526 (SCLC-P), NCI-H211 (SCLC-P), NCI-H510 (non-SCLC-P), and NCI-H1963 (SCLC-P) cell lines were transduced with either non-targeting CRISPR sgRNA or C11orf53 specific sgRNAs for four days. The cell viability was determined by cell counting assay, n=3, two-tailed unpaired Student’s *t* test. \*\**P* < 0.01; \**P* < 0.05. F) NCI-H526 and NCI-H211 cell lines were transduced with either non-targeting CRISPR sgRNA or C11orf53 specific sgRNAs for four days. The cell morphology was shown under bright field. G) and H) The protein levels of C11orf53, Cyclin B1, Cyclin E1, Cyclin A2, Cleaved PARP, Cleaved Caspase 3, and Cleaved Caspase 7 were determined by western blot in NCI-H526 (G) and NCI-H211 (H) cell lines with C11orf53 CRISPR depletion. HSP90 was used as internal control, n=2.

The *C11orf53* gene encodes for a 288-amino acid protein product without any known functional domain/motif or cellular localization signal. When analyzing the co-dependency scores of top genes that are selectively essential in SCLC-P subtype, POU2F3 and C11orf53 have an overall correlation coefficient of 0.812 in the SCLC-P subtype and 0.357 of all 17 SCLC cell lines (Fig. 2B). This prompted us to investigate the functional relationship between C11orf53 and POU2F3 in P-subtype SCLC. In published SCLC patient samples (EGAS00001000925), we have detected a positive correlation between the expression levels of POU2F3 and C11orf53 (Fig. 2C). Therefore, this gene may function as a new biomarker for SCLC-P subtype cells.

To study the function of C11orf53 in SCLC cells, we generated our homemade polyclonal antibody for C11orf53 and detected the protein levels of C11orf53 in four different SCLC cell lines. As shown in Figure 2D, we have detected very high levels of C11orf53 protein in the NCI-H526 and NCI-H211 SCLC-P subtype cell lines, however there were no detectable signals from NCI-H510 nor NCI-H1963 non-SCLC-P subtype cell lines. Depletion of C11orf53 in SCLC cells dramatically reduced cell viability (Fig. 2E) and induced a significant change of the cell morphology (Fig. 2F). To further determine how this gene contributes to cell viability, we determined the protein levels of several factors involved in both cell cycle regulatory checkpoints and apoptosis from two different SCLC-P cell lines. As shown in Figure 2G and 2H, in both SCLC-P cell lines we found depletion of *C11orf53* via CRISPR knockout could induce cleavage of PARP, Caspase-3, and Caspase-7. Our results suggest that the *C11orf53* gene is critical for the maintenance of SCLC-P cell viability.

### C11orf53 regulates lineage-specific genes expression at broad enhancer domains

To understand how C11orf53 globally impacts gene expression in SCLC cells, we conducted RNA-seq in NCI-H526 cell lines transduced with either non-targeting CRISPR sgRNA, or two distinct C11orf53 specific sgRNAs. As shown in Figure 3A, we found a total of 3,719 and 3,259 genes that were significantly down- and up-regulated, respectively, upon *C11orf53* depletion. Moreover, GSEA pathway analysis have identified several key cell growth biological processes, such as Myc and E2F signaling pathway-dependent gene signatures, which were dramatically downregulated upon *C11orf53* depletion (Fig. 3B). To understand how this small protein regulates gene expression, we first sought to determine the cellular localization of C11orf53 by cell fractionation assay. Surprisingly, we found the C11orf53 protein could be detected in the cytosol, soluble nuclear, and chromatin (insoluble nuclear) fractions (Fig. 3C), even though it obviously has no nuclear localization signal (NLS). To further investigate how this protein regulates gene expression at chromatin, we conducted ChIP-seq with two of our home-made polyclonal antibodies. Based on our ChIP-seq analysis, both of our antibodies show consistent performance, and the peaks are highly correlated (Fig. S3A, B). In general, we have detected 8820 specific C11orf53 peaks in NCI-H526 cells (Fig. 3D). Interestingly, we found that most of these peaks were localized at intergenic or intron regions, indicating that C11orf53 may regulate gene expression at distal enhancer elements. In addition, the motif analysis has identified POU2F3 motif being the top matched hit (p-value 1e-3053) motif as C11orf53 peaks (Fig. 3E). This result implies that there might be a similar functionality or genetic interaction between POU2F3 and C11orf53 within SCLC-P subtype cells.

**Figure 3.**
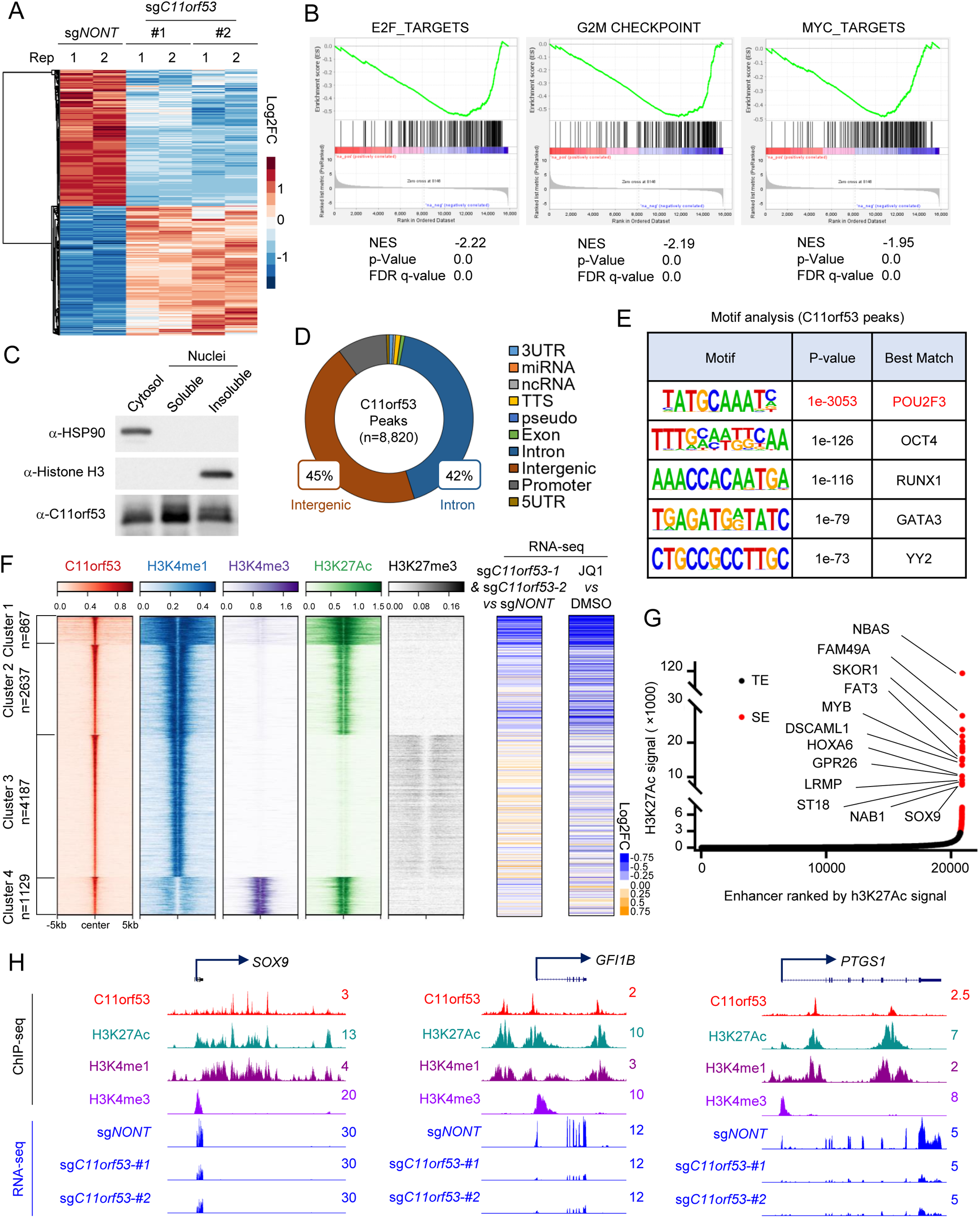
C11orf53 regulates lineage-specific genes expression at broad enhancer domains. A) RNA-seq was conducted with NCI-H526 cell lines transduced with either non-targeting CRISPR sgRNA or C11orf53 specific sgRNAs for four days. The heat map shows the differentially regulated genes, n=2. B) The GSEA plot shows the enrichment of E2F, G2M checkpoint, and MYC pathways genes enriched in the downregulated genes with C11orf53 depletion. C) The protein levels of C11orf53 in cytosol, soluble nuclear fraction, and insoluble nuclear fraction was determined by western blot. HSP90 was used as cytoplasmic protein control, and the histone H3 was used as nuclear insoluble protein control, n=2. D) The pie plot shows the annotation and distribution of C11orf53 peaks at the genome. E) Motif analysis shows the best matched motifs occupied by C11orf53. F) The total C11orf53 peaks were divided into four clusters based on kmeans clustering. The histone marks were further centered on C11orf53 peaks in each cluster (left). RNA-seq was conducted with cells transduced with either non-targeting CRISPR sgRNA or C11orf53 specific sgRNAs. The log2 foldchange heat maps shows the expression change of nearest genes to C11orf53 peaks (middle), n=2. RNA-seq was conducted with cells treated with either DMSO or JQ1 (1 μM). The log2 foldchange heat maps shows the expression change of nearest genes to C11ORF53 peaks (right), n=2. G) The Histone H3 lysine 27 acetylation (H3K27ac) signals from chromatin immunoprecipitation (ChIP) sequencing identifies putative super enhancers (SEs) in NCI-H526 cells. Hockey-stick plot representing the normalized rank and signals of H3K27Ac. Representative SE-associated genes that are occupied by C11orf53 are labeled. H) Representative tracks showing the enhancer binding of C11orf53, which contributes to activation of gene expression.

To further characterize the function of C11orf53 at the genome-wide scale and understand how C11orf53 regulates gene expression at distal enhancers, we divided all C11orf53 peaks into four clusters based on k-means clustering. As shown in Figure 3F, we have detected a very significant overlap between enhancer marker H3K4me1 and C11orf53 peaks. Interestingly, both of Cluster 1 and 2 peaks are also enriched with active enhancer marker H3K27ac, suggesting that C11orf53 may function as a transcriptional activator at active enhancers. We further integrated our RNA-seq data with the ChIP-seq data and identified the expression change of the genes that are nearest to C11orf53 peaks. As shown in the middle panel of Figure 3F and Figure S3C, we found that genes nearest to Cluster 1 peaks are dramatically reduced upon *C11orf53* depletion, where a very broad H3K4me1 peaks were detected. Indeed, as shown in Figure 3G, most of the super-enhancer (SE) associated genes (labeled) are reduced after C11orf53 depletion. In general, around 66% (71/108) SE-associated genes are occupied by C11orf53, in which 73.2% (52/71) genes were down-regulated upon C11orf53 depletion (Fig. S3D). Pathway analysis with the genes that are nearest to Cluster 1 peaks show a significant enrichment in several neuronal function and differentiation pathways (Fig. S3E). Next, to further confirm the role of C11orf53 involvement in the regulation of SE-associated genes, we determined the gene expression profiles within cells treated with either DMSO or JQ1. As shown in the right panel of Figure 3F, Figure 3G, S3F-I, the JQ1 treatment has induced a very similar change in the gene expression profile as *C11orf53* depletion. Notably, multiple tuft cell-specific genes were down-regulated after *C11orf53* depletion or JQ1 treatment, in accordance with POU2F3 function^12^ (Fig. 3H, S3J).

### Loss of C11orf53 reduces enhancer activity and chromatin accessibility

Since C11orf53 occupies active enhancers and is critical for maintenance of the expression levels of the enhancer-nearby genes, we asked whether loss of C11orf53 impacts the protein levels of enhancer histone marks, such as H3K4me1 and H3K27ac. We conducted western blot in cells transduced with either non-targeting CRISPR sgRNA or two distinct C11orf53 sgRNAs. As a result, we found that loss of *C11orf53* expression does not affect the bulk of histone modifications (Fig. 4A). Next, we conducted ChIP-seq to determine whether there were loci-specific changes of active enhancer marks (H3K4me1/H3K27ac). As shown in Figure 4B and 4C, *C11orf53* depletion strongly reduced H3K27ac levels and a moderate reduction in H3K4me1 levels. Then we centered the log2 fold-change of H3K4me1/H3K27ac levels at the previously defined four C11orf53-specifc clusters in Figure 3F. As shown in Figure 4D, we have detected a very strong reduction of H3K27ac and H3K4me1 histone marks near C11orf53 peaks. Consistently, there is also a dramatic reduction of super enhancer signals in C11orf53-depleted cells (Fig. 4E, F). Finally, we conducted ATAC-seq experiments to determine whether the chromatin accessibility has been altered in C11orf53-depleted cells. As a result, we have detected a total of 55,111 ATAC-seq peaks with 7,805 of those peaks being occupied by C11orf53 (Fig. 4G). We further separated the total ATAC-seq peaks into two groups, depending on whether they were co-bound with C11orf53. As shown in Figure 4H, we found that Group 1 peaks (ATAC-seq+/C11orf53+) were strongly reduced upon both sgRNAs treatment (Fig. 4H, left panel, 4I, and S4A, B), while Group 2 peaks (ATAC-seq+/C11orf53-) were not significantly reduced upon *C11orf53* depletion (Fig. 4H, right panel, Fig. S4C). In summary, our results have shown a novel function of C11orf53 in mediating chromatin accessibility and maintaining enhancer activity in SCLC-P subtype cells.

**Figure 4.**
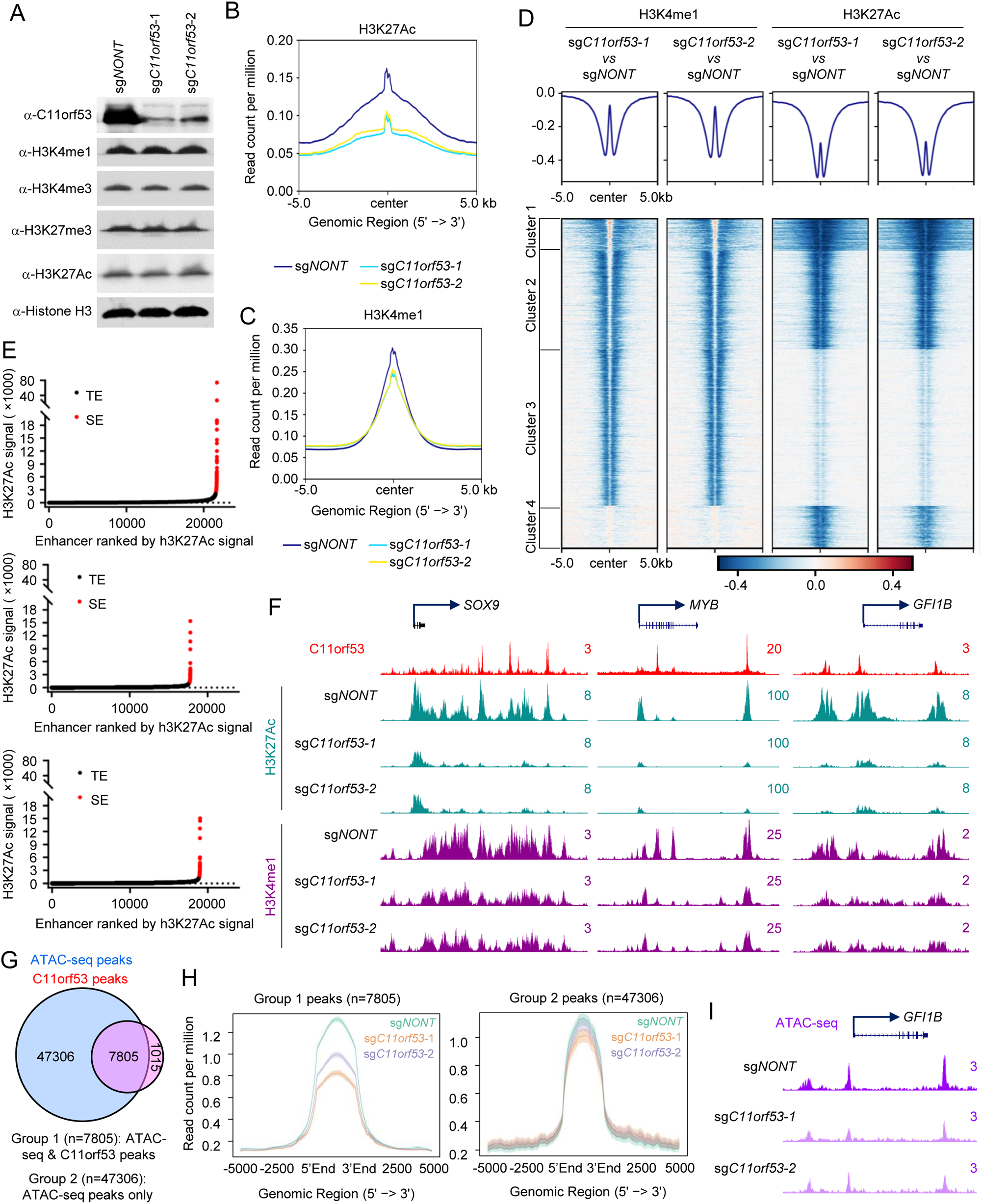
Loss of C11orf53 reduces enhancer activity and chromatin accessibility. A) NCI-H526 cell lines transduced with either non-targeting CRISPR sgRNA or C11orf53 specific sgRNAs for four days. The protein levels of C11orf53, H3K4me1, H3K4me3, H3K27me3, and H3K27Ac levels were determined by western blot. Total histone H3 was used as internal control, n=2. The average plot shows the total H3K27Ac (B), and H3K4me1 (C) peaks in NCI-H526 cell lines transduced with either non-targeting CRISPR sgRNA or C11orf53 specific sgRNAs. Chromatin from drosophila S2 cells (10%) was used as spike-in. D) The log2 fold change heatmap shows the loss of H3K4me1 and H3K27Ac signal at C11ORF53 peaks after C11ORF53 depletion. E) Histone H3K27ac signals from chromatin immunoprecipitation (ChIP) sequencing identifies putative super enhancers (SEs) in NCI-H526 cells transduced with either non-targeting CRISPR sgRNA or C11orf53 specific sgRNAs. Hockey-stick plot representing the normalized rank and signals of H3K27ac. Representative of top ranked SE-associated genes from each group are labeled. F) Representative track examples have shown the H3K27ac levels at SOX9, GFI1B, and PTGS1 gene loci in cells transduced with either CRISPR sgRNA or C11orf53 specific sgRNAs. G) The venn-diagram shows the overlap between ATAC-seq peaks and C11orf53 peaks in NCI-H526 cells. H) The average plot shows the ATAC-seq signal between cells transduced with either CRISPR sgRNA or C11orf53 specific sgRNAs. The Group 1 peaks are ATAC-seq/C11orf53 common peaks. Group 2 peaks are ATAC-seq alone peaks. I) Representative tracks showing ATAC-seq peaks at *GFI1B* gene loci in cells transduced with either CRISPR sgRNA or C11orf53 specific sgRNAs.

### C11orf53 is a co-activator of POU2F3 in SCLC cells

As an uncharacterized 288-amino acid protein, there is no obvious functional domain/motif within C11orf53. To study the underlying molecular basis of how C11orf53 is recruited to chromatin and regulates gene expression, we purified GFP-tagged proteins from SCLC cell line NCI-H526 which stably express either GFP or GFP-tagged C11orf53. By mass spectrometry analysis, we found POU2F3 as one of the top candidates that interact with C11orf53 (Fig. 5A, B). We further performed immunoprecipitation experiments with whole cell lysate from NCI-H526 cells to confirm the protein-protein interaction between the endogenous C11orf53 and POU2F3 (Fig. 5C, D). To understand how C11orf53 interacts with POU2F3, we truncated POU2F3 into different truncated fragments (Fig. 5E) and purified each of these GFP-tagged fragments in HEK293T cells co-transfected with Halo-tagged C11orf53 (Fig. 5F). As shown in Figure 5G, we found that the POU domain within the POU2F3 protein is critical for the interaction with C11orf53. Since C11orf53 protein has no DNA or histone binding domain, we hypothesized that it may be transported into the nuclei and recruited to chromatin by POU2F3. To test this hypothesis, we transfected HEK293T cells (which do not express either POU2F3 or C11orf53) with Halo-tag or Halo-tagged POU2F3 in the presence of GFP-tagged C11orf53. As shown in Figure S5A, the chromatin bound C11orf53 was dramatically increased in the presence of POU2F3. To confirm this observation in SCLC cells, we depleted POU2F3 by CRIPSR knockout in NCI-H526 cells for two days and determined the chromatin-bound C11orf53 levels. As shown in Figure S5B, depletion of POU2F3 could also reduce the chromatin bound C11orf53. These results imply that C11orf53 might function as a co-activator that is recruited to chromatin by POU2F3^16^.

**Figure 5.**
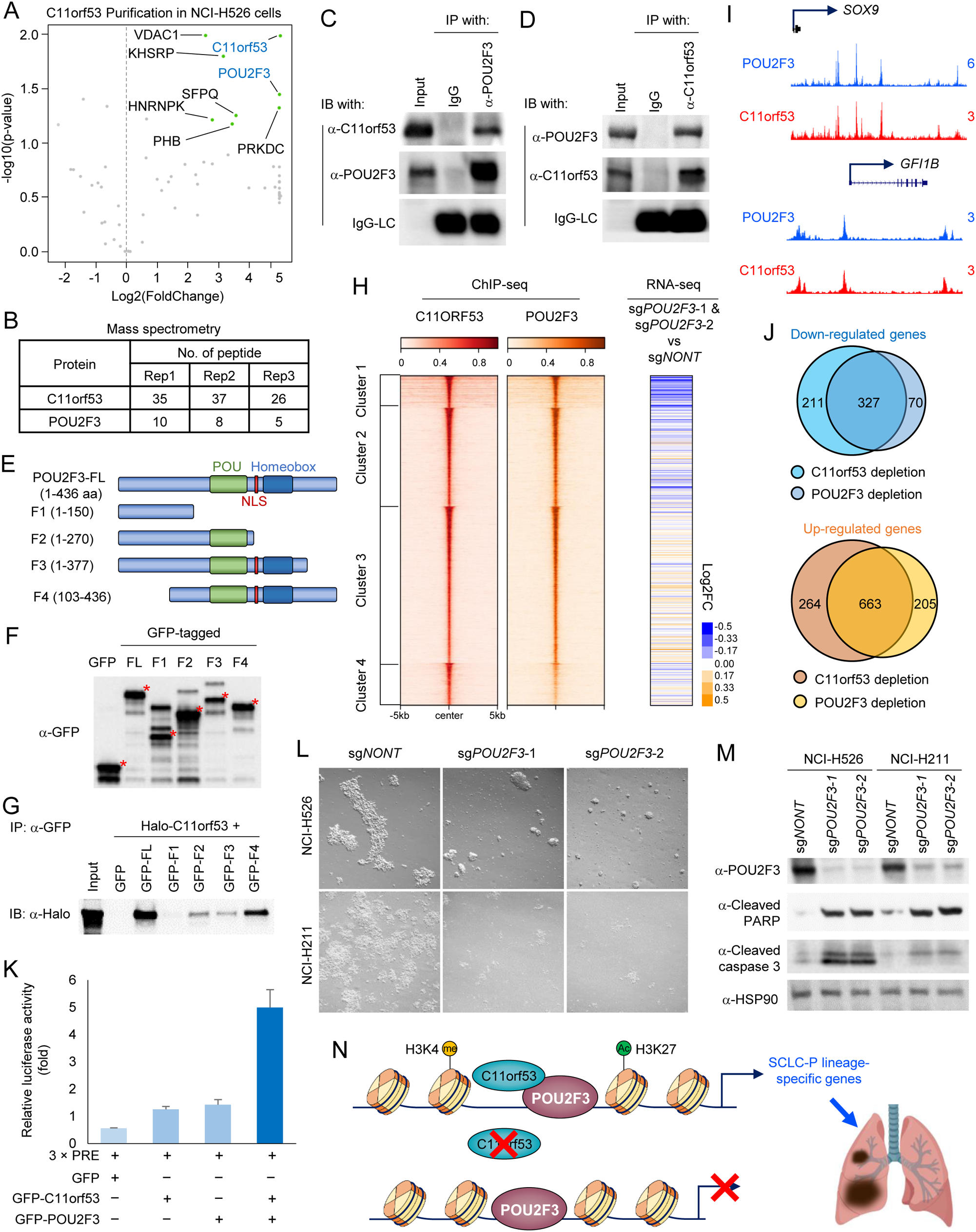
C11orf53 is a co-activator of POU2F3 in SCLC cells. NCI-H526 cells were infected by lenti-virus expressing either GFP or GFP-tagged C11orf53. The GFP-tagged proteins were purified and subjected to mass spectrometry analysis. A) The volcano plot shows the significant enriched protein during GFP-tagged C11orf53 purification, n=3. Peptide numbers of POU2F3 and the bait protein C11orf53 were shown (B). IP of endogenous POU2F3 from NCI-H526 cells followed by IB for POU2F3 and C11orf53 (C) and vice versa (D). IgG was used as a negative control, n=2. E) Schematic diagram depicting the domain organization of the human POU2F3 protein. The indicated fragments were sub-cloned as a GFP-tag fusion into pLJM-GFP plasmid. F) Whole-cell lysates were used for western blot with GFP antibodies in cells transfected with empty vector (GFP) or different POU2F3 fragments in (E), n=2. G) Whole-cell lysates were used for immunoprecipitation (IP) with GFP antibodies followed by immunoblotting (IB) for Halo-tag in cells co-transfected with Halo-tagged C11orf53, and either empty vector (GFP) or POU2F3 fragments in (E), n=2. H) Sorted and centered heatmaps generated from ChIP-seq data show the occupancy of C11orf53 and POU2F3 in NCI-H526 SCLC cells. All rows are centered on C11orf53 four clusters as divided by kmeans clustering (left). RNA-seq was conducted with cells transduced with either non-targeting CRISPR sgRNA or POU2F3 specific sgRNAs. The log2 foldchange heat maps shows the expression change of nearest genes to C11orf53 peaks (middle), n=2 (right). I) The representative tracks show the co-localization of POU2F3 and C11orf53 at *SOX9* and *GFI1B* gene loci. J) The venn-diagram shows the overlap of targeted genes between POU2F3 and C11orf53. K) HEK293T cells were transfected with GFP-tagged C11orf53 and/or POU2F3 in the presence of 3×PRE reporter plasmid. Luciferase assay was performed 48 hours after transfection, n=3. L) NCI-H526 and NCI-H211 cell lines were transduced with either non-targeting CRISPR sgRNA or POU2F3 specific sgRNAs for four days. The cell morphology was shown under bright field. M) The protein levels of POU2F3, Cleaved PARP, and Cleaved Caspase 3 were determined by western blot. HSP90 was used as internal control, n=2. N) Model of C11orf53 functions as co-activator of POU2F3 in SCLC cells.

Finally, to determine a potential co-function between C11orf53 and POU2F3, we conducted ChIP-seq to determine the chromatin occupancy of POU2F3 in NCI-H526 cells. As we expected, there is a significant overlap between POU2F3 and C11orf53 occupancy across the genome (Fig. 5H, left panel, 5I), and *POU2F3* depletion also reduced the expression of Group 1 clustered genes, which is similar compared to *C11orf53* depletion (Fig. 5H, right panel). In total, there are 327 genes that are co-downregulated by C11orf53 or POU2F3 depletion (fold change > 2), in which we have identified numerous tuft cell lineage genes—which has been demonstrated to be targeted genes of POU2F3 (Fig. 5J, S5C). Since the chromatin binding motif between C11orf53 and POU2F3 is extremely similar, we have introduced luciferase assay to further confirm the co-function between these two factors. As shown in Figure 5K, we found co-transfection of C11orf53 and POU2F3 has the strongest effect on inducing luciferase expression. Indeed, similar to *C11orf53* depletion, loss of POU2F3 also dramatically induces apoptosis and reduces cell viability in two different SCLC-P cell lines (Fig. 5L, M). In summary, our model has shown that this previous uncharacterized protein, C11orf53, functions as a co-activator of POU2F3 and maintains chromatin accessibility at POU2F3-targeted genes in SCLC cells (Fig. 5N).

## Discussion

The POU family proteins are a large class of DNA-binding transcription factors that are divided into six classes, with POU2F3 belonging to the POU II class, which includes POU2F1 (Oct-1) and POU2F2 (Oct-2)^17^. As the only known binding partner of POU II class, POU2AF1 (POU Class 2 Homeobox Associating Factor 1) functions as coactivator of POU2F1, and to a lesser extent than POU2F1, as a coactivator of POU2F2^18-21^. Our current study has provided evidence for a previously uncharacterized protein, C11orf53, functioning as a co-activator of POU2F3 (also known as Oct-11). Therefore, we are defining this gene based on its novel function and classifying it as the Second POU Class 2 Homeobox Associating Factor (or *POU2AF2*). Based on our mass spectrometry analysis of purified C11orf53, we did not detect POU2F1 or POU2F2, which are both expressed in NCI-H526 cells based on our RNA-seq data. This result suggested that C11orf53 is quite a unique co-activator of POU2F3. Indeed, it is known that POU2AF1 binds to POU2F1 and POU2F2 using its alpha-helical segment, while C11orf53 does not display a similar structure^22^. Therefore, it would be interesting to determine the crystal structure of C11orf53 bounded to POU2F3, but also challenging because of the disordered nature of C11orf53 structure.

The function of POU2F3 in small-cell lung cancer has been identified by genome-wide CRISPR screening in previous studies^12^. Therefore, it has been widely used as the biomarker that defines SCLC-P subtype, which expresses markers of the chemosensory lineage instead of neuroendocrine markers. Mechanistically, POU2F3 binds to distal enhancer elements and drives the expression of the lineage-specific genes. Interestingly, as a non-histone/DNA binding co-activator, C11orf53 is recruited to chromatin by the DNA binding transcription factor, POU2F3, and maintains openness of the chromatin to regulate the expression of POU2F3-targeted genes. Based on our model (Fig. 5N), C11orf53 may not function as a canonical pioneer factor that recruits transcription factor to chromatin. Therefore, it is critical to identify SCLC-specific pioneer factors that mediates POU2F3 recruitment to enhancers.

Interestingly, based on our genome-wide RNA-seq analysis, there are approximately 32% of C11orf53-targeted genes that are not affected upon *POU2F3* depletion. It is possible that the sgRNA for C11orf53 is more efficient than that for POU2F3. However, it is also possible that C11orf53 may have additional functions other than co-activators of POU2F3 based on our cellular fractionation assays. Future studies may be focused on the cytoplasmic function of C11orf53 in SCLC cells.

Dysregulation or mutations within enhancer binding factors have been identified as direct drives for many types of cancers^23,24^. Our current studies have shown a critical role of C11orf53 at broad active enhancer domains. Consequently, loss of C11orf53 leads to a loci-specific reduction of H3K27ac levels and gene expression, which is similar to the effects of JQ1 treatment. Our result is consistent with previous reports that SCLC cells are more sensitive to BET inhibitor treatment^11,25,26^. Finally, our study sheds light on the potential impact of targeting C11orf53 or C11orf53/POU2F3 heterodimer for new therapeutic approaches against SCLC-P subtype due to its specificity and high dependency in this particular cancer type. Therefore, establishing small-molecule inhibitors or peptide drugs that can disrupt C11orf53/POU2F3 interaction may specifically inhibit POU2F3-dependent transcriptional programming in SCLC cells and thus aid in the development of a more personalized approach to SCLC therapy.

## Materials and Methods

### Antibodies and Reagents

POU2F3 (#36135), H3K27ac (#8173S), H3K4me1 (#5326S), H3K4me3 (#9751), H3K27me3 (#9733), histone H3 (#4499S), Cyclin B1 (#12231), Cyclin E1 (#20808), Cyclin A2 (#91500), Cleaved-PARP (#5625), Cleaved Caspase 3 (#9664) and Cleaved-Caspase 7 (#8438) antibodies were purchased from Cell Signaling. Tubulin antibody (E7) was purchased from Developmental Studies Hybridoma Bank. HSP90 (sc-7947) and GFP (sc-9996) antibodies were purchased from Santa Cruz. Halo-tag (G9211) antibody was purchased from Promega. The C11orf53 antibody was produced in rabbit in house by using full-length C11orf53 recombinant protein as antigen.

### Cell Lines

HEK293T cells were obtained from ATCC, and then maintained with DMEM (Gibco, Gaithersburg, MD) containing 10% FBS (Sigma). The SCLC cell lines NCI-H526, NCI-H211, NCI0H510, and NCI-H1963 were obtained from ATCC, and were maintained with ATCC-formulated RPMI-1640 medium containing 10% FBS (Sigma). The *Drosophila* S2 cells were maintained in HyClone™ SFX-Insect Cell Culture Media containing 10% FBS (Sigma).

### Immunoprecipitation (IP)

The IP experiment was performed as described before. Briefly, the cells were lysed in the lysis buffer (50mM Tris pH 8.0, 150 mM NaCl, 0.5% Triton X100, 10% Glycerol, protease inhibitors, and benzonase). After centrifugation at max speed at 4°C for 15 min, the supernatants were collected and incubated with the primary antibody and immobilized Protein A/G (Santa Cruz) at 4°C overnight with rotation. Then the samples were washed with lysis buffer four times and boiled in 5× SDS sample loading buffer.

### RNA Interference, CRISPR-mediated knockouts, and Real-time PCR

Designed sgRNAs were cloned into lentiCRISPR v2 (Addgene, 52961) vector. lentiviral-mediated CRISPR/Cas9 knockout was described previously. Oligo sequences used in this manuscript were as follows: sgNONT (GCTGAAAAAGGAAGGAGTTGA), sgC11orf53-1 (GTGACGTCTACACCTCCAGCG), sgC11orf53-2 (GAGAGGCAACTCGTGCTGGG), sgPOU2F3-1 (GCCCACGCTTAGGGAGATGTG), and sgPOU2F3-2 (GTCCTACCAAATACTTCACTG),

### RNA-seq and analysis

RNA-seq was conducted as previously described. Paramagnetic beads coupled with oligo d(T) are combined with total RNA to isolate poly(A)+ transcripts based on NEBNext® Poly(A) mRNA Magnetic Isolation Module manual. All remaining steps for library construction were used according to the manufacturer’s recommendations. Samples were pooled and sequenced on a HiSeq with a read length configuration of 150 PE. Gene counts were computed by HTSeq and used as an input for edgeR 3.0.85257. Genes with Benjamini-Hochburg adjusted p-values less than 0.01 were considered to be differentially expressed (unless otherwise specified).

### ChIP-seq Assay and analysis

ChIP-seq was performed as described previously. For histone modifications, 10% of Drosophila chromatin was used as spike-in control. For ChIP-seq analysis, all the peaks were called with the MACS v2.1.0 software using default parameters and corresponding input samples. Metaplots and heatmaps were generated using ngsplot database to display ChIP-seq signals. Peak annotation, motif analysis, and super enhancer analysis were performed with HOMER and ChIPseeker. Pathway analysis was performed with Metascape and ChIPseeker.

### ATAC-seq and analysis

ATAC-seq was performed as described previously. In brief, frozen cells were thawed and washed once with PBS and then resuspended in 500 uL of cold ATAC lysis buffer. The cell number was assessed by Cellometer Auto 2000 (Nexcelom Bioscience). 50K to 100K nuclei were then centrifuge (pre-chilled) at 500 g for 10 min. Supernatant was then removed and the nuclei were resuspended in 50 uL of tagmentated DNA by pipetting up and down six times. The reactions were incubated at 37 °C for 30 min in a thermomixer shaking at 1,000 rpm, and then cleaned up by the MiniElute reaction clean up kit (Qiagen). Tagmentated DNA was amplified with barcode primers. Library quality and quantity were assessed with Qubit 2.0 DNA HS Assay (ThermoFisher, Massachusetts, USA), Tapestation High Sensitivity D1000 Assay (Agilent Technologies, California, USA), and QuantStudio® 5 System (Applied Biosystems, California, USA). Equimolar pooling of libraries was performed based on QC values and sequenced on an Illumina® HiSeq (Illumina, California, USA) with a read length configuration of 150 PE for 50M PE reads (25M in each direction) per sample. ATAC-seq reads are shifted + 4 bp and − 5 bp for positive and negative strands respectively using alignmentSieve function from deepTools package. ATAC-seq peaks are called with Macs v2.1.0. sgNONT ATAC-seq peaks were intersected with C11orf53 peaks using bedtools.

### Mass Spectrometry Sample Preparation and analysis

Mass spectrometry was performed as described previously. Protein pellet was denatured in 50 μL of 8 M Urea/0.4 M Ammonium Bicarbonate followed by reduction in 2 μL of 100 mM DTT. The digests were acidified to 0.5% trifluoroacetic acid (TFA) and the peptides were then desalted on C18 Sep-Paks (Waters). The pooled extracts were dried in a vacuum concentrator and resuspended in 30 uL of 5% ACN/0.1% FA for LC-MS analysis. Peptides were analyzed by LC-MS/MS using a Dionex UltiMate 3000 Rapid Separation LC (RSLC) system and a linear ion trap-Orbitrap hybrid Elite mass spectrometer (Thermo Fisher Scientific Inc, San Jose, CA).

### Statistical Analyses

For statistical analyses, GraphPad Prism 7, Microsoft Excel, and R were used. All data involving a statistical analysis being reported that met the criteria to use the appropriate statistical tests; for the normal distribution of data, the empirical rule was used to infer the distribution. For growth curves and time-course, RNA-seq t-tests were calculated between the area-under-the-curve (AUC) values. Statistical tests used are reported in the figure legends.

## Data availability

NGS data generated for this study are available upon request

## Acknowledgements

We would like to thank Dr. Feng Zhang for the kind gifts of the Px330 and lentiCRISPR v2 vectors.

## Authors’ contributions

LW and ZZ designed the study. NT, APS, and LW performed all the biochemistry and sequencing experiments. ZZ performed the bioinformatics analysis. NT, APS, LW wrote the manuscript. All authors read and approved the final manuscript.

## Supplementary Figure

**Figure S1.**
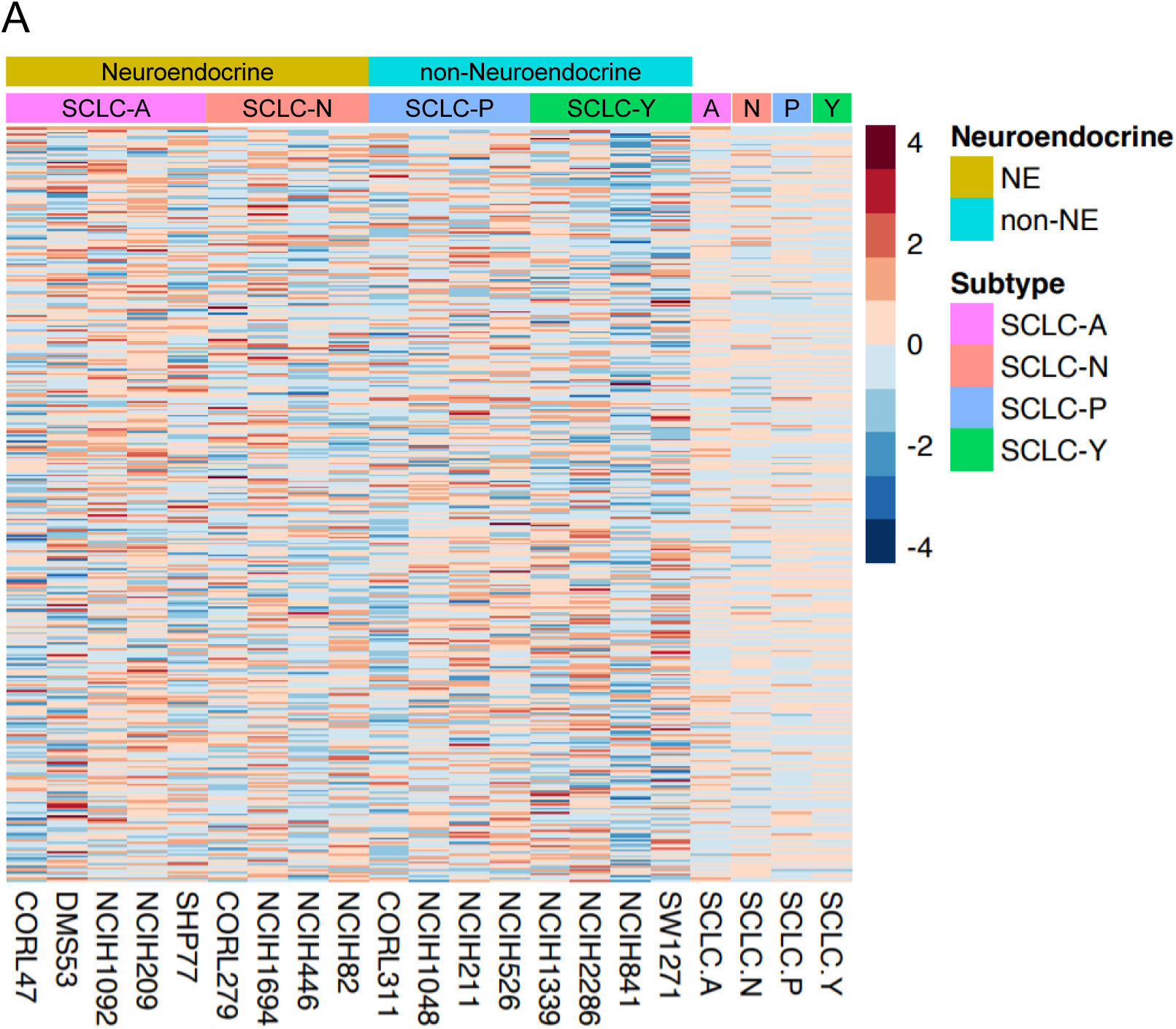
Landscape of SCLC subtype-specific dependency. A) RNAseq TPM gene expression data for the 17 cell lines were retrieved from DepMap Public 21Q3 datasets. The Z-score heatmap showed the gene expression in the same order as Figure 1A.

**Figure S2.**
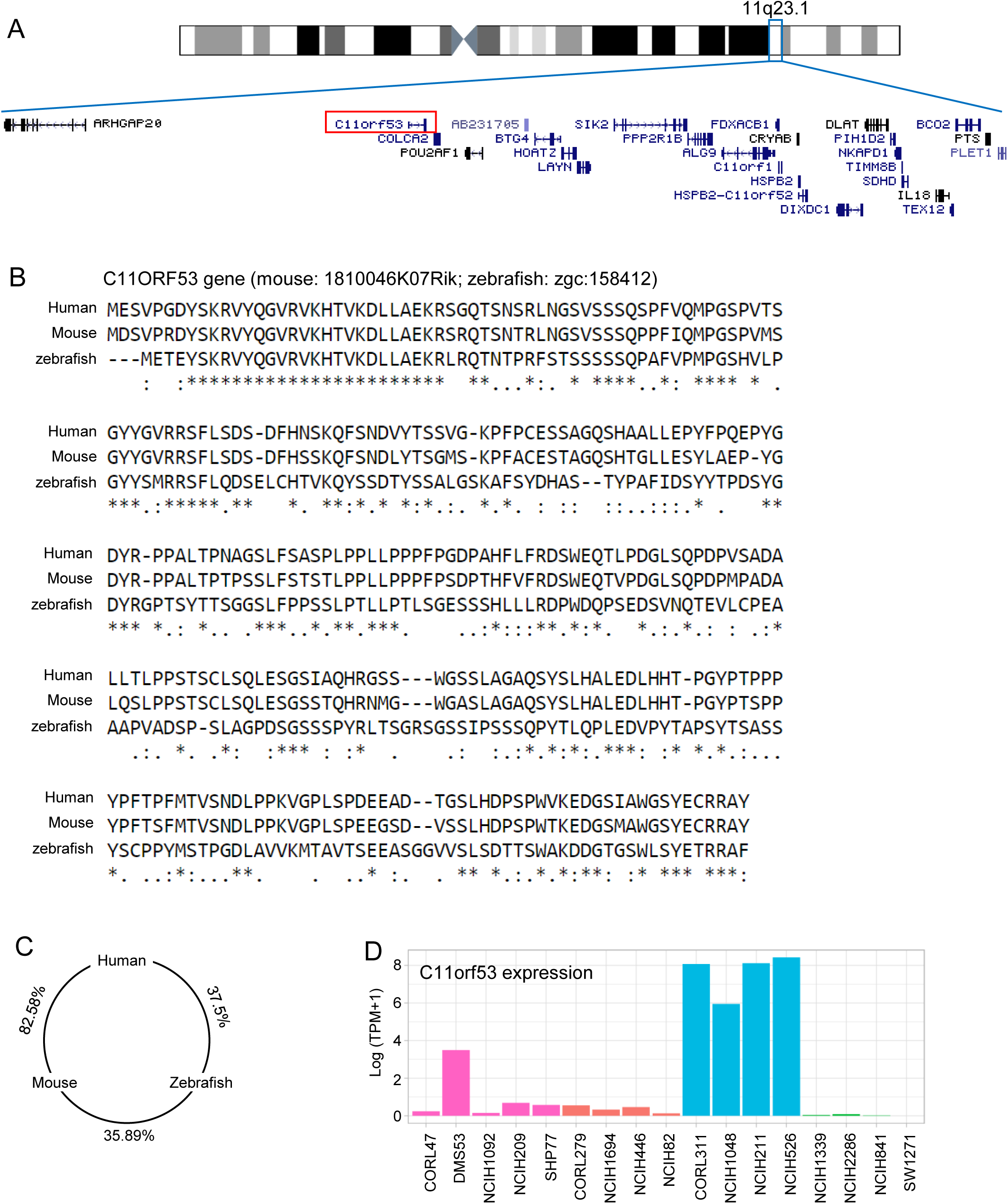
Identification of C11orf53 as a new SCLC-P subtype marker. A) The location of C11orf53 gene on chromosome 11q23.1. B) Comparison of protein produce of C11orf53 gene (1810046K07Rik in mouse and zgc: 158412 in zebrafish) between human, mouse and zebrafish by CLUSTALW. C) The aligned score for similarity between human C11orf53, mouse, and zebrafish zgc: 158412 was calculated by ClustalW sequence alignment. D) C11orf53 expression in 17 SCLC cell lines. Log (TPM+1) expression data were retrieved from DepMap Public 21Q3 dataset.

**Figure S3.**
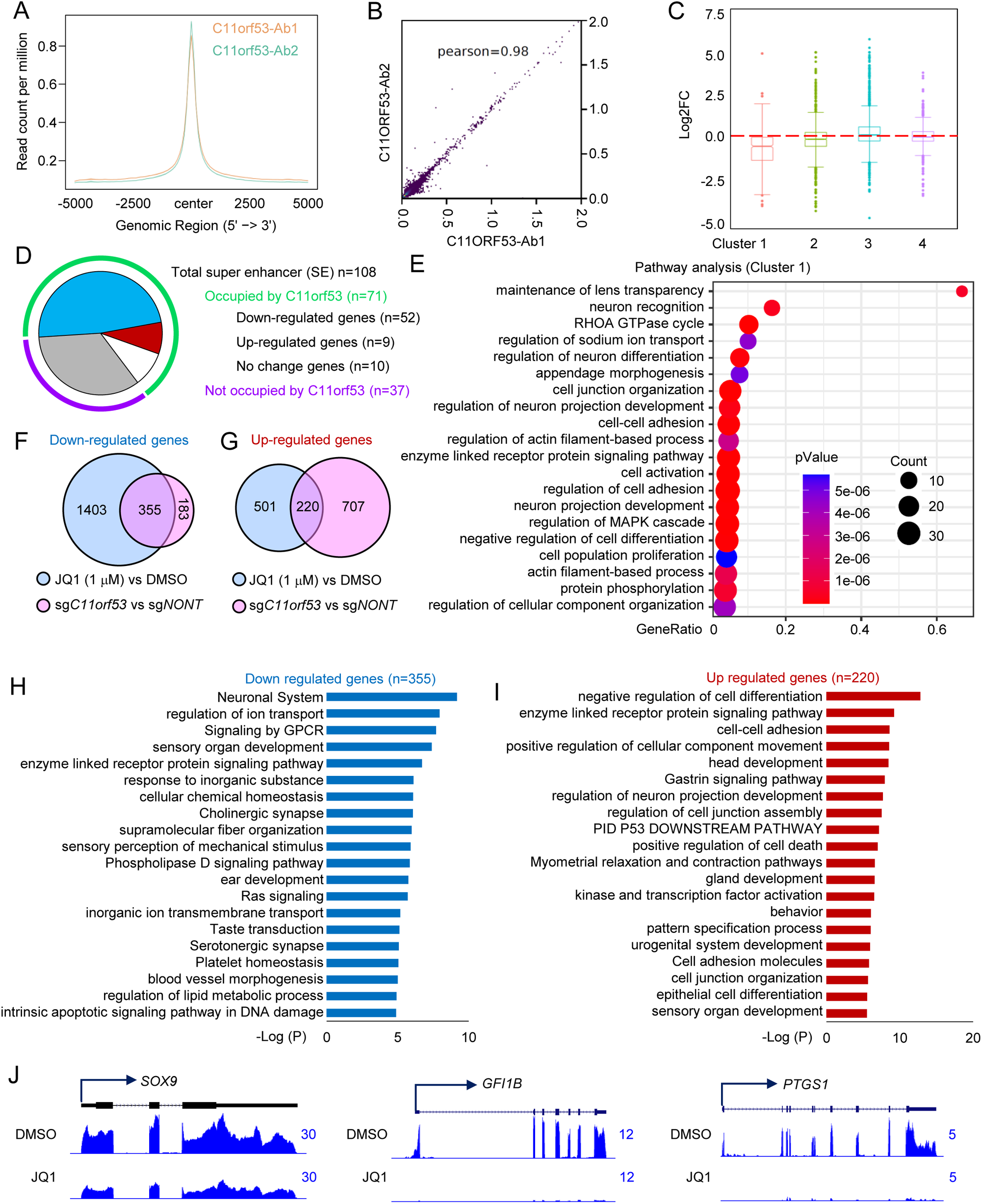
C11orf53 regulates lineage-specific genes expression at broad enhancer domains. The average plot shows the co-localization of C11orf53 peaks detected by two antibodies (A), and the Pearson Correlation coefficient value was determined by using multiBigwigSummary and plotCorrelation functions from deepTools (B). C) The box plot shows the expression change of genes nearest to C11orf53 peaks in each of the four clusters. D) The pie plot shows the impact of C11orf53 on super enhancer (SE) associated genes in NCI-H526 cells. E) Pathways analysis with genes nearest to Cluster 1 peaks of C11orf53. F) and G) The Venn-diagram shows the overlap of down-regulated gene (F) and up-regulated genes (G) in NCI-H526 cells treated with JQ1 (1 μM) or two distinct C11orf53 sgRNAs, n=2. Pathway analysis by Metascape of 355 genes (Log2FC > 1) that are co-downregulated by JQ1 treatment or C11orf53 depletion (H), and 220 genes (Log2FC > 1) that are co-upregulated by JQ1 treatment or C11orf53 depletion (I). J) The representative tracks show the treatment of JQ1 also reduced the expression levels of C11orf53 targeted genes.

**Figure S4.**
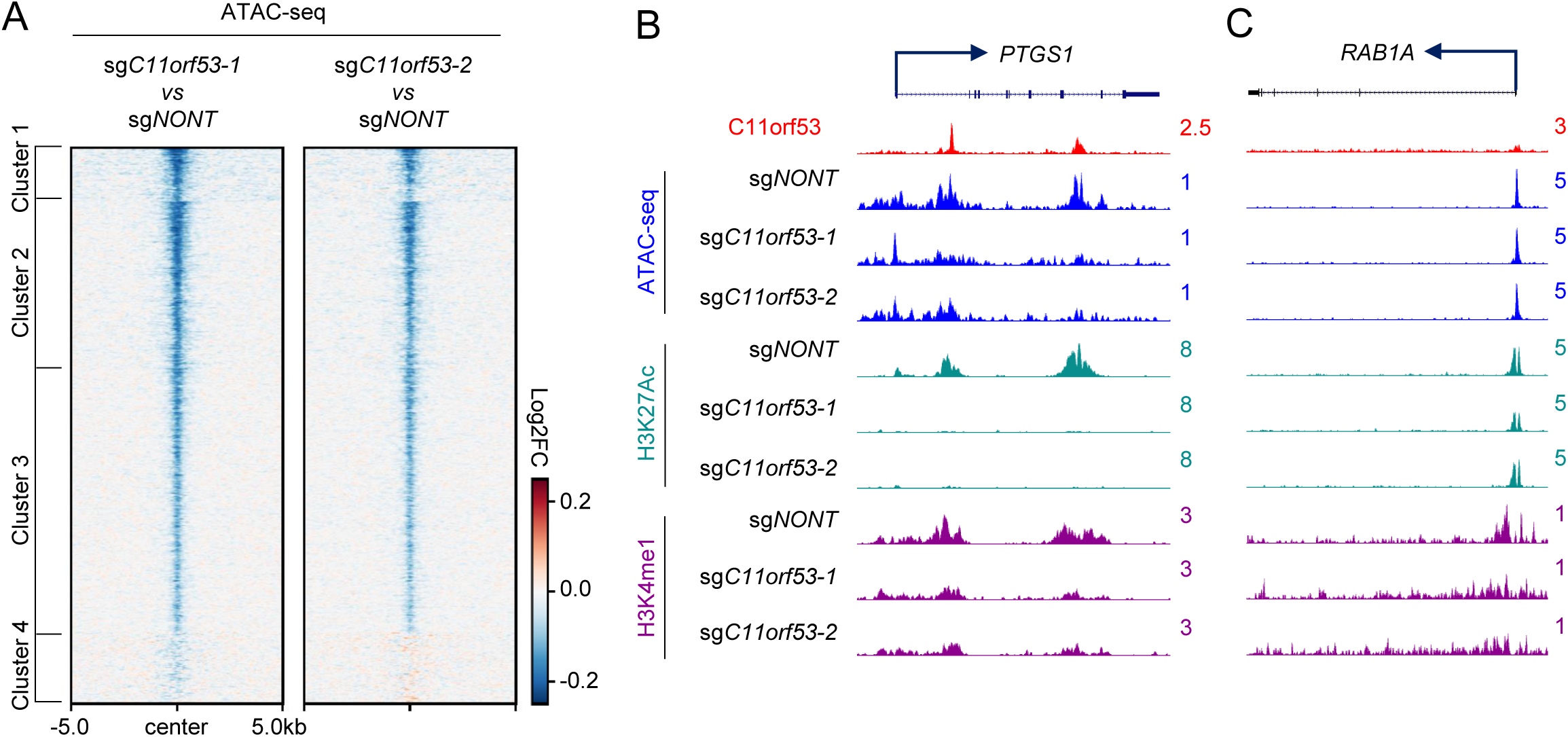
Loss of C11orf53 reduces enhancer activity and chromatin accessibility. A) The ATAC-seq was performed in NCI-H526 cells transduced with either non-targeting CRISPR sgRNA, or two distinct C11orf53 specific sgRNAs. The log2FC change of ATAC-seq signal was centered on C11orf53 peaks. The representative tracks show the ATAC-seq signal and active enhancer marks at PTGS1 gene loci (C11ORF53 binding) (B), and RAB1A gene loci (no C11orf53 binding) (C).

**Figure S5.**
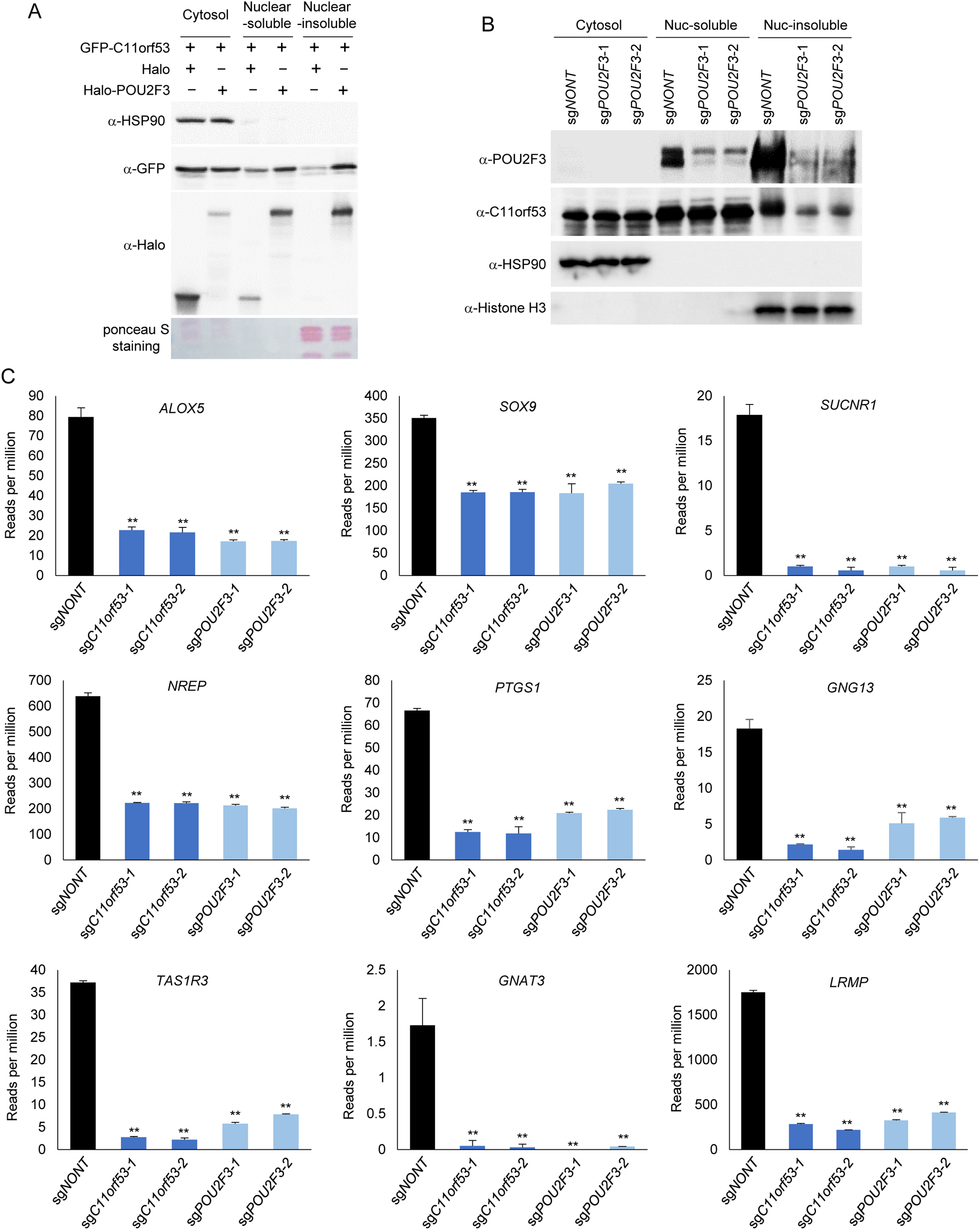
C11orf53 is recruited by POU2F3 to chromatin. A) HEK293T cells were transfected with GFP-tagged C11orf53 in the presence of either Halo-tag, or Halo-tagged POU2F3. The cellular fractionation was isolated and the protein levels of C11orf53 and POU2F3 was determined by GFP or Halo-tag antibodies. HSP90 was used as cytoplasmic protein control, and the histone H3 was used as nuclear insoluble protein control, n=2. B) The SCLC cell line NCI-H526 was transduced with either non-targeting CRISPR sgRNA, or two distinct POU2F3 specific sgRNAs for 72 hours, followed by cellular fractionation assay. The protein levels of POU2F3 and C11orf53 in each fraction was determined by western blot. HSP90 was used as cytoplasmic protein control, and the histone H3 was used as nuclear insoluble protein control, n=2. C) The SCLC cell line NCI-H526 was transduced with either non-targeting CRISPR sgRNA, or two distinct C11orf53 specific sgRNAs, or two distinct POU2F3 specific sgRNAs for four days, respectively. The mRNA levels of tuft cell specific markers, including ALOX5, SOX9, SUCNR1, NREP, PTGS1, GNG13, TAS1R3, GNAT3, and LRMP in each group was determined by RNA-seq, n=2, two-tailed unpaired Student’s *t* test. \*\**P* < 0.01; \**P* < 0.05.

## Reference

1 Sung, H. et al. Global cancer statistics 2020: GLOBOCAN estimates of incidence and mortality worldwide for 36 cancers in 185 countries. Ca-Cancer J Clin 71, 209–249, doi:10.3322/caac.21660 (2021).

2 Gardner, E. E. et al. Chemosensitive Relapse in Small Cell Lung Cancer Proceeds through an EZH2-SLFN11 Axis. Cancer Cell 31, 286–299, doi:10.1016/j.ccell.2017.01.006 (2017).

3 Gazdar, A. F., Bunn, P. A. & Minna, J. D. Small-cell lung cancer: what we know, what we need to know and the path forward. Nat Rev Cancer 17, 725–737, doi:10.1038/nrc.2017.87 (2017).

4 Rudin, C. M., Brambilla, E., Faivre-Finn, C. & Sage, J. Small-cell lung cancer. Nat Rev Dis Primers 7, doi:ARTN 3 10.1038/s41572-020-00235-0 (2021).

5 Miller, K. D. et al. Cancer treatment and survivorship statistics, 2019. Ca-Cancer J Clin 69, 363–385, doi:10.3322/caac.21565 (2019).

6 Rudin, C. M. et al. Molecular subtypes of small cell lung cancer: a synthesis of human and mouse model data (vol 19, pg 289, 2019). Nat Rev Cancer 19, 415–415, doi:10.1038/s41568-019-0164-2 (2019).

7 Zhang, W. et al. Small cell lung cancer tumors and preclinical models display heterogeneity of neuroendocrine phenotypes. Transl Lung Cancer R 7, 32-+, doi:10.21037/tlcr.2018.02.02 (2018).

8 Yang, D. et al. Intertumoral Heterogeneity in SCLC Is Influenced by the Cell Type of Origin. Cancer Discov 8, 1316–1331, doi:10.1158/2159-8290.Cd-17-0987 (2018).

9 Schwendenwein, A. et al. Molecular profiles of small cell lung cancer subtypes: Therapeutic implications. Mol Ther-Oncolytics 20, 470–483, doi:10.1016/j.omto.2021.02.004 (2021).

10 Bebber, C. M. et al. Ferroptosis response segregates small cell lung cancer (SCLC) neuroendocrine subtypes. Nat Commun 12, doi:ARTN 2048 10.1038/s41467-021-22336-4 (2021).

11 Szczepanski, A. P. et al. ASXL3 bridges BRD4 to BAP1 complex and governs enhancer activity in small cell lung cancer. Genome Med 12, doi:ARTN 63 10.1186/s13073-020-00760-3 (2020).

12 Huang, Y. H. et al. POU2F3 is a master regulator of a tuft cell-like variant of small cell lung cancer. Gene Dev 32, 915–928, doi:10.1101/gad.314815.118 (2018).

13 Shai, R. et al. Gene expression profiling identifies molecular subtypes of gliomas (vol 22, pg 4918, 2003). Oncogene 25, 4256–4256, doi:10.1038/sj.onc.1209746 (2006).

14 Bertucci, F. et al. Gene expression profiling identifies molecular subtypes of inflammatory breast cancer. Cancer Res 65, 2170–2178, doi:Doi 10.1158/0008-5472.Can-04-4115 (2005).

15 Tlemsani, C. et al. SCLC-CellMiner: A Resource for Small Cell Lung Cancer Cell Line Genomics and Pharmacology Based on Genomic Signatures. Cell Rep 33, doi:ARTN 108296 10.1016/j.celrep.2020.108296 (2020).

16 Moritsugu, R. et al. Functional analysis of the nuclear localization signal of the POU transcription factor Skn-1a in epidermal keratinocytes. Int J Mol Med 34, 539–544, doi:10.3892/ijmm.2014.1803 (2014).

17 Ryan, A. K. & Rosenfeld, M. G. POU domain family values: Flexibility, partnerships, and developmental codes. Gene Dev 11, 1207–1225, doi:DOI 10.1101/gad.11.10.1207 (1997).

18 Gstaiger, M., Knoepfel, L., Georgiev, O., Schaffner, W. & Hovens, C. M. A B-Cell Coactivator of Octamer-Binding Transcription Factors. Nature 373, 360–362, doi:DOI 10.1038/373360a0 (1995).

19 Strubin, M., Newell, J. W. & Matthias, P. Obf-1, a Novel B-Cell-Specific Coactivator That Stimulates Immunoglobulin Promoter Activity through Association with Octamer-Binding Proteins. Cell 80, 497–506, doi:Doi 10.1016/0092-8674(95)90500-6 (1995).

20 Shore, P., Dietrich, W. & Corcoran, L. M. Oct-2 regulates CD36 gene expression via a consensus octamer, which excludes the co-activator OBF-1. Nucleic Acids Res 30, 1767–1773, doi:DOI 10.1093/nar/30.8.1767 (2002).

21 Shakya, A. et al. Oct1 and OCA-B are selectively required for CD4 memory T cell function. J Exp Med 212, 2115–2131, doi:10.1084/jem.20150363 (2015).

22 Chasman, D., Cepek, K., Sharp, P. A. & Pabo, C. O. Crystal structure of an OCA-B peptide bound to an Oct-1 POU domain/octamer DNA complex: specific recognition of a protein-DNA interface. Gene Dev 13, 2650–2657, doi:DOI 10.1101/gad.13.20.2650 (1999).

23 Plass, C. et al. Mutations in regulators of the epigenome and their connections to global chromatin patterns in cancer. Nat Rev Genet 14, 765–780, doi:10.1038/nrg3554 (2013).

24 Khan, P. et al. Epigenetic landscape of small cell lung cancer: small image of a giant recalcitrant disease. Semin Cancer Biol, doi:10.1016/j.semcancer.2020.11.006 (2020).

25 Lenhart, R. et al. Sensitivity of Small Cell Lung Cancer to BET Inhibition Is Mediated by Regulation of ASCL1 Gene Expression. Mol Cancer Ther 14, 2167–2174, doi:10.1158/1535-7163.Mct-15-0037 (2015).

26 Lam, L. T. et al. Vulnerability of Small-Cell Lung Cancer to Apoptosis Induced by the Combination of BET Bromodomain Proteins and BCL2 Inhibitors. Mol Cancer Ther 16, 1511–1520, doi:10.1158/1535-7163.Mct-16-0459 (2017).

